# Chlorotonils exhibit potent activity against *Mycobacterium tuberculosis*, while resistance is mediated by MmpR5-MmpL5

**DOI:** 10.64898/2025.12.09.693342

**Authors:** Felix Deschner, Melissa D Chengalroyen, Lauren Ames, Diana Quach, Rodrigo Aguilera Olvera, Barbara Bosch, Anthony Castro, Heather Kim, Kaushik Raman, Natalie Thornton, Joshua Wallach, Fernanda Rodrigues da Costa, Renee Allen, Andréanne Lupien, Mandisa Zuma, Sasha Lynch, Joe Pogliano, Joseph Sugie, Jeremy M Rock, Dirk Schnappinger, Tanya Parish, Valerie Mizrahi, Michael DeJesus, Rolf Müller, Jennifer Herrmann

## Abstract

Treatment of *Mycobacterium tuberculosis* (Mtb) is challenging and requires administration of at least four different antibiotics. Unfortunately, multi-drug resistant Mtb strains continue to emerge, undermining the effectiveness of current treatment regimens and highlighting the urgent need for new therapeutics. In this study, we evaluated the potential of natural product-derived chlorotonils as anti-Mtb agents. We demonstrate that chlorotonils exhibit nanomolar potency against a range of attenuated and virulent Mtb strains. Mechanistic studies and resistance profiling in Mtb revealed that chlorotonils affect both lipid and energy metabolism. Through systems biology approaches, including the construction of an Mtb CRISPRi library specifically designed for chemical-genomic profiling, we identified MmpR5/MmpL5 as major driver of chlorotonil-resistance in Mtb leading also to cross-resistance with bedaquiline. Our findings highlight chlorotonils as valuable chemical tools to further dissect the role and function of the MmpS5-MmpL5 efflux pump in drug-resistant Mtb.

## 1. Introduction

Tuberculosis (TB), a life-threatening disease caused by *Mycobacterium tuberculosis* (Mtb), remains a leading cause of death from infectious disease worldwide. It is estimated that a quarter of the global population has been infected with Mtb^1^. While TB most commonly affects the lungs - resulting in pulmonary TB - it can also present as extra-pulmonary TB, impacting various parts of the body. These manifestations include gastrointestinal TB, meningeal TB, pericardial TB, skeletal TB, and lymph node TB^2^. Other clinically significant mycobacterial pathogens include *Mycobacterium abscessus complex* and *Mycobacterium avium complex*, which together account for more than 90% of all documented cases of non-tuberculous mycobacteria pulmonary disease (NTM-PD)^3^.

Standard TB treatment for drug-susceptible TB is complex and requires a combination of several antibiotics - typically rifampicin, isoniazid, pyrazinamide and ethambutol - administered for a minimum of four to six months^4^. Treating rifampicin mono-resistant (RR), multidrug-resistant (MDR) and extensively drug-resistant (XDR) TB can be even more challenging, where treatment can extend up to 20 months. However, significant progress has been made in shortening the treatment period for patients with RR- or MDR-TB by the introduction of *e.g*., the all-oral 6-month BPaLM regime composed of bedaquiline (BDQ), pretonamide, linezolid, and moxifloxacin^1,4–6^. Although the ATP synthase inhibitor BDQ was only approved in 2012 specifically for MDR and XDR-TB, resistance against BDQ has already been reported, further complicating efforts to control the disease^7,8^. In recognition of the growing threat posed by drug-resistant Mtb, the World Health Organization (WHO) updated its priority pathogen list in 2024. Rifampicin-resistant Mtb is now classified as a Priority 1 (critical) pathogen - alongside MDR Gram-negative bacteria - underscoring the urgent need for new and more effective treatment options^9,10^.

Natural products remain a major source of new antimicrobial drugs^11^. Chlorotonils are macrocyclic antibiotics originally isolated from the myxobacterium *Sorangium cellulosum*, and exhibiting potent activity against MDR pathogens such as vancomycin-resistant *Enterococcus faecium* (VRE) or methicillin-resistant *Staphylococcus aureus* (MRSA), as well as the malaria-causing parasite *Plasmodium falciparum*^12–14^. In *S. aureus*, chlorotonils display activity against MDR pathogens through a complex mode of action (MoA) that involves targeting bacterial membrane lipids, the peptidoglycan synthesis protein YbjG, and the methionine aminopeptidase, ultimately leading to rapid depolarization of the bacterial membrane as their primary activity^15^. Notably, resistance development in *S. aureus* occurs only after prolonged exposure to sub-lethal concentrations, with resistance to the compound achieved through increased FarE-mediated lipid efflux.

Here we sought to characterize the activity of chlorotonils against Mtb. We quantified inhibition of growth and killing of Mtb by chlorotonils, performed a functional genomics screen utilizing an Mtb CRISPRi pooled library and Bacterial Cytological Profiling (BCP), and generated resistant mutants, to explore their MoA and potential for the treatment of TB.

## 2. Results

### Chlorotonils are active against Mtb

We performed standard antimicrobial susceptibility testing of the natural product Chlorotonil A (ChA) and the two semi-synthetic derivatives Chlorotonil B1-Epo2 (ChB1-Epo2; first series frontrunner^11^) and dehalogenil (DHG; lead^12^) against attenuated Mtb H37Ra. Excellent activity was observed for all three derivatives with minimal inhibitory concentrations (MICs) in the range of 0.1-0.2 µg/mL (**Table 1**), in line with previous results found for Gram-positive bacteria^13,14^. We then tested the compounds against the virulent Mtb H37Rv, as well as *M. smegmatis*, and the NTM strains *M. avium* and *M. abscessus*. H37Rv showed similar susceptibility to the natural product derivatives when compared to the attenuated H37Ra strain (MICs 0.1-1 µg/mL), whereas DHG exhibited no activity against the NTM strains under the conditions tested.

**Table 1:** Activity profiling of chlorotonils against Mycobacteria.

| Strain | MIC [µg/mL] |  |  |
| --- | --- | --- | --- |
|  | DHG | ChA | ChB1-Epo2 |
| *Mycobacterium tuberculosis* H37Ra | 0.125 | 0.1-0.2 | 0.125 |
| *Mycobacterium tuberculosis* H37Rv | 0.136-0.3 | 0.6 | 1.25 |
| *Mycobacterium avium* subsp. *hominissuis* MAH104 | >40 | >1 | >10 |
| *Mycobacterium abscessus* ATCC19977 | >40 | >1 | >10 |
| *Mycobacterium smegmatis* ATCC700084 (mc<sup>2</sup>155) | >8* | >8 | >8 |

\*as part of mechanistic studies, inhibitory activity against *M. smegmatis* was confirmed according to assay setup at higher concentrations.

### DHG shows a slow onset of activity in Mtb without inducing membrane depolarization

Having found significant anti-tubercular activity for chlorotonils, we next determined their impact on the viability of Mtb. In previous studies focusing on *S. aureus*, we demonstrated that chlorotonils act rapidly, inducing a viable but non-culturable (VBNC) state immediately upon exposure^15,16^. We therefore conducted an analogous assay using Mtb H37Ra exposed to 0.1 µg/mL or 1 µg/mL of DHG, corresponding to ∼0.8x and ∼8x the MIC, or DMSO as the control. At the indicated time points, samples were plated to perform a viable cell count (**Figure 1**A). At 0.1 µg/mL, a bacteriostatic-type effect was observed for the first six days, followed by a slight increase in colony-forming unit (CFU) count until day eight, plateauing at roughly 10% of the CFU count observed for the DMSO control. At the higher drug concentration (1 µg/mL), a slow but steady decline in CFU counts was observed throughout the 11-day treatment period leading to a ∼2 log_10_ reduction in CFU compared to the inoculum. While a slower onset of drug activity in Mtb compared to fast-growing Gram-positive bacteria is generally expected, our results indicate that DHG does not induce an immediate viable but nonculturable (VNBC) state in Mtb as previously reported for *S. aureus*^17^, suggesting species-dependent differences in DHG’s mechanism of action.

**Figure 1:**
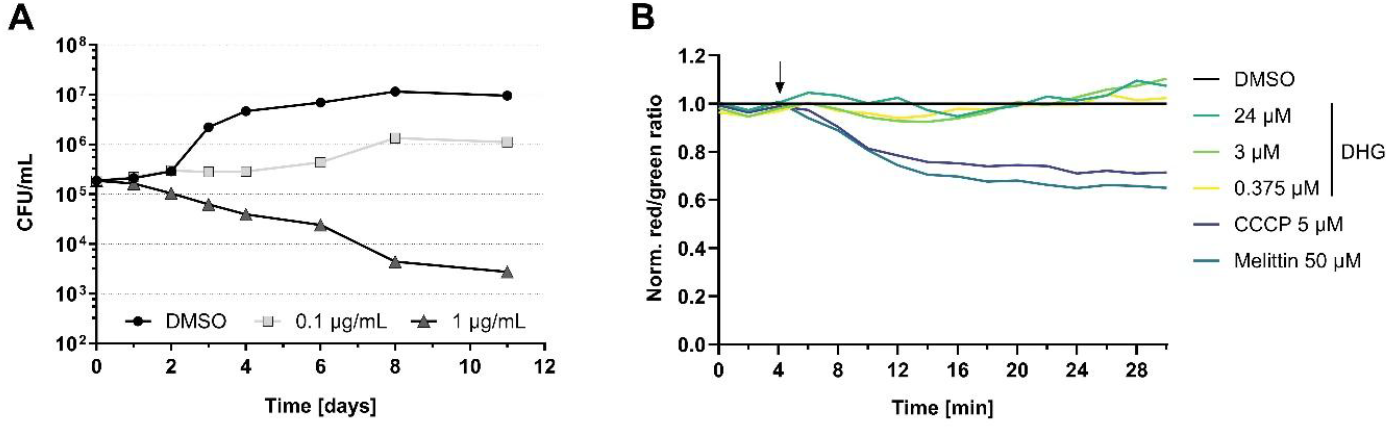
DHG acts slowly against M. tuberculosis and does not induce membrane depolarization. A) Viable cell count of Mtb H37Ra after DHG (or DMSO) treatment. Samples were drawn at indicated time points and plates were incubated at 37°C for four weeks. B) DiOC_2_(3) assay does not indicate membrane depolarization of Mtb cells upon treatment with DHG for 30 minutes. Arrow indicates compound addition. Red/green signal ratio was normalized against DMSO; experiment was performed in two technical replicates with the line representing the average control level.

Given its different onset of action in different bacteria, we assessed chlorotonil’s reported membrane depolarization mechanism, previously described in *S. aureus*^15^, in Mtb using a standard DiOC_2_(3) assay (3,3’-Diethyloxacarbocyanine Iodide, **Figure 1**B). Mtb H37Ra was treated for 30 minutes with different concentrations DHG, the positive controls 5 µM Carbonyl cyanide m-chlorophenyl hydrazone (CCCP) and 50 µM Melittin, while 1% DMSO served as negative control for signal normalization. Consistent with the rather slow decline in CFUs, DHG did not induce membrane depolarization within 30 minutes, whereas both CCCP and Melittin induced significant effects, confirming a different activity profile of chlorotonils in Mtb compared to *S. aureus*^17^.

To further explore the MoA of chlorotonils in Mtb, we performed Myco-Bacterial Cytological Profiling (Myco-BCP) using the Linnaeus platform (Linnaeus Bioscience Inc., USA)^18^ (**Figure 1**A/B, **Table SI 1**). This platform applies quantitative bacterial cell biology powered by automated image analysis to predict the mode of action of novel antimicrobials^18^. Currently, it can discriminate between inhibition of nine different pathways covering ATP synthase and proton motive force (ATP/PMF), cell wall (CW1-2; peptidoglycan or arabinogalactan/MmpL3, respectively), DNA, fatty acid and mycolic acid (FA/MA), membrane (MEMB), transcription (TRSC) and translation (TRSL)^18^. Treatment with DHG resulted in slightly elongated cells, a minor increase in cell permeability at higher treatment concentrations and heterogeneous, patchy membrane staining, including discrete bright foci indicative of localized membrane accumulation (**Figure 2**A). Hence, we found very high similarity scores of the DHG activity profile with ATP and proton motive force inhibitors BDQ and CCCP (similarity scores were 87 and 86, respectively). Myco-BCP groups BDQ and CCCP together based on the morphology that results from treatment after 48 or 120 hours. While CCCP is a classical ionophore that directly collapses membrane potential, BDQ inhibits ATP synthase and depletes ATP without immediate membrane depolarization^19,20^. The similarity in cytological profiles therefore reflects convergence on impaired energy production similar to BDQ, rather than an ionophore-like activity as observed for CCCP.

**Figure 2:**
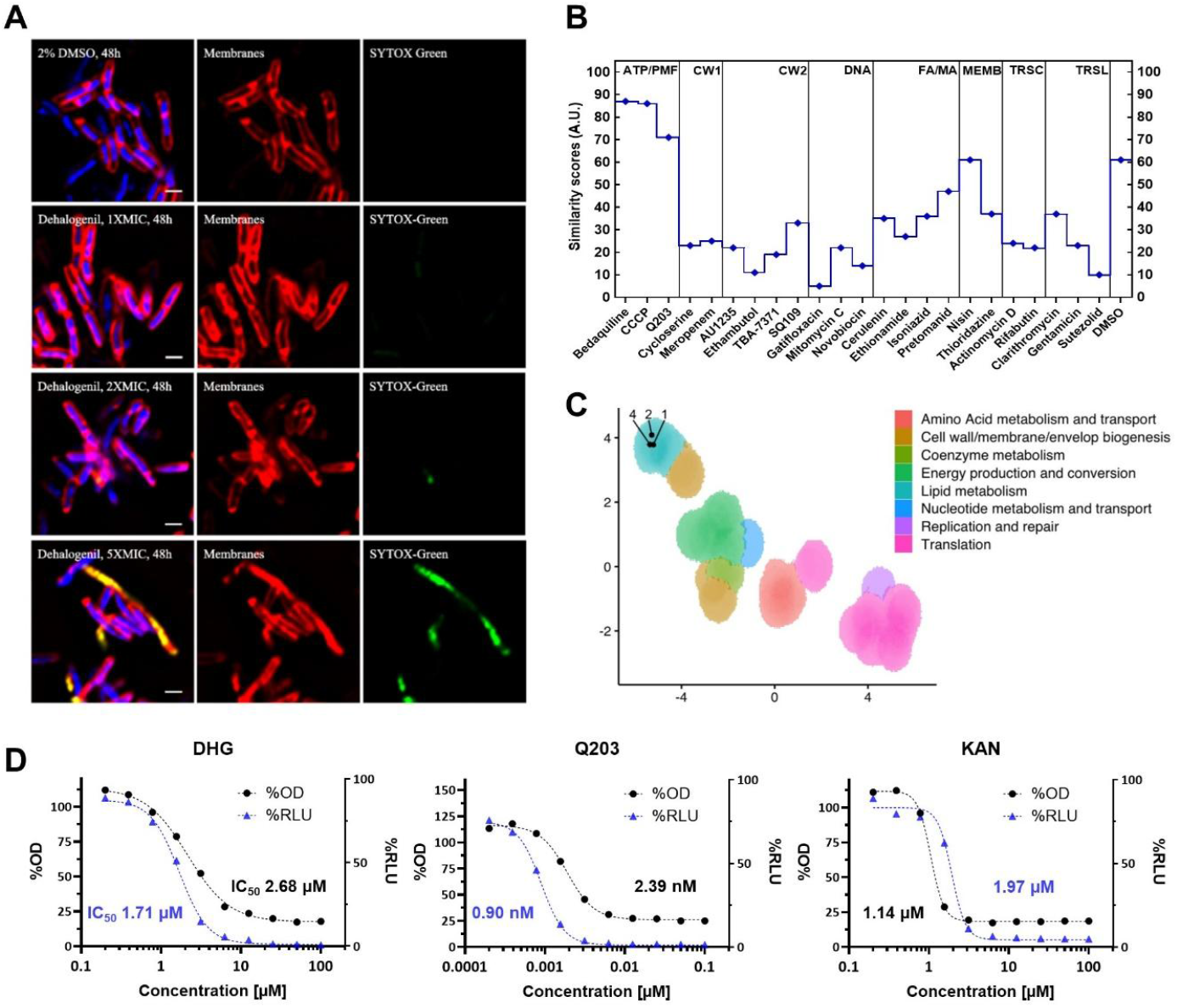
Evidence implicating energy and lipid metabolism in the mechanism of antimycobacterial activity of chlorotonils. A) Mycobacterial cytological profiling (Linnaeus; Myco-BCP) with corresponding similarity score in B. Scale bar is 1 µM. Pathways: ATP synthase and proton motive force (ATP/PMF), cell wall (CW1-2; peptidoglycan or arabinogalactan/MmpL3, respectively), DNA, fatty acid and mycolic acid (FA/MA), membrane (MEMB), transcription (TRSC), translation (TRSL) and DMSO. C) Morphological profiling of M. smegmatis in response to treatment with DHG (1x, 2x, and 4x MIC; MIC 0.25 mM) illustrates that bacillary morphotypes cluster in UMAP space with CRISPRi hypomorphs in genes involved in lipid metabolism. Black circle, untreated. D) Mtb H37Rv-LP was exposed to compounds and ATP levels were measured after 24 h by the addition of BacTiter-Glo reagent and reading luminescence (RLU). OD_590_ was measured after 5 days. Q203 was used as a positive control (ATP depletion), and kanamycin (KAN) was used as a negative control (no effect). Data was normalized against control cells.

In a parallel approach, complementary to Myco-BCP, we also performed a morphological profiling of *M. smegmatis* treated with DHG by generating quantitative imaging of the drug-treated cells for comparison against an arrayed *M. smegmatis* CRISPRi hypomorph library (**Figure 2**C)^21^. Exposure of wildtype *M. smegmatis* to increasing concentrations of DHG (1x, 2x, and 4x MIC) resulted in bacillary morphotypes that clustered very tightly in UMAP space in an area defined by hypomorphs in lipid metabolism genes, and proximal to those observed for *wag31, accA3, fas, mmpL3* and *accD4/5* hypomorphs. This result suggested an association between lipid metabolism and the MoA of chlorotonil in *M. smegmatis*.

Given that chlorotonils share phenotypic and mechanistic similarities with compounds acting within the cytoplasmic membrane of Mtb, while not causing membrane depolarization, we hypothesized that chlorotonils might interfere with ATP production and energy-related pathways in a BDQ-like manner. However, we only found mild ATP depletion in DHG-treated cells (**Figure 2**D) using a BacTiter-Glo assay and by comparing IC_50_ values derived from luminescence (ATP content) and OD_590_ data. The ATP-based IC_50_ was approximately 36% lower than the growth-based IC_50_ in DHG-treated cells, while this effect was more pronounced (62%) for cells treated with the positive control drug, Q203, which targets the QcrB subunit of the cytochrome *bc*_1_-*aa*_3_ complex^22^.

### Functional genomics reveals silencing of *mmpR5* as key resistance factor and links DHG activity to lipid metabolism

To further investigate the MoA of chlorotonils in Mtb, we turned to genome-wide CRISPRi profiling, a technique that provides a genome-wide approach to identify genes that determine susceptibility of Mtb to antitubercular compounds^23^. However, our current CRISPRi library in Mtb consists of 96,700 sgRNA, which limits access to this profiling approach due to the high cost associated with sequencing such a large library. To overcome this limitation and identify the genes determining intrinsic resistance of Mtb to growth inhibition by DHG, we designed and constructed a new CRISPRi library specifically for chemical-genomic studies consisting of 31,143 sgRNAs, which reduces the sequencing costs of chemical-genomic profiling experiments by ∼67% while still enabling tuneable knockdown of gene expression for genes essential for *in vitro* growth.

Five of the genes whose silencing increased sensitivity of Mtb to several concentrations of DHG (*rv0204c, rv0479c, rv1109c, rv1707* and *rv3802c*) encode membrane proteins mostly of unknown function, which may be related to the membrane-targeting activity of DHG (**Figure 3**A). Importantly, the sensitizing genes included *mmpL5* and *mmpS5*, which encode a drug efflux pump known to export BDQ and other drugs. Transcription of *mmpL5* and *mmpS5* is normally repressed by the transcriptional regulator MmpR5^24^ and as such, silencing of *mmpR5* likely increased resistance of Mtb to DHG via upregulation of *mmpL5* and *mmpS5*. Other notable genes that increased resistance to DHG upon silencing included *acpM, kasA, kasB, fabD, accA3, fas*, and *accD*, all of which are involved in mycolic acid synthesis. Thus, the chemical-genomic profiling identified genes in lipid metabolism as both sensitizers and resistors to DHG.

**Figure 3:**
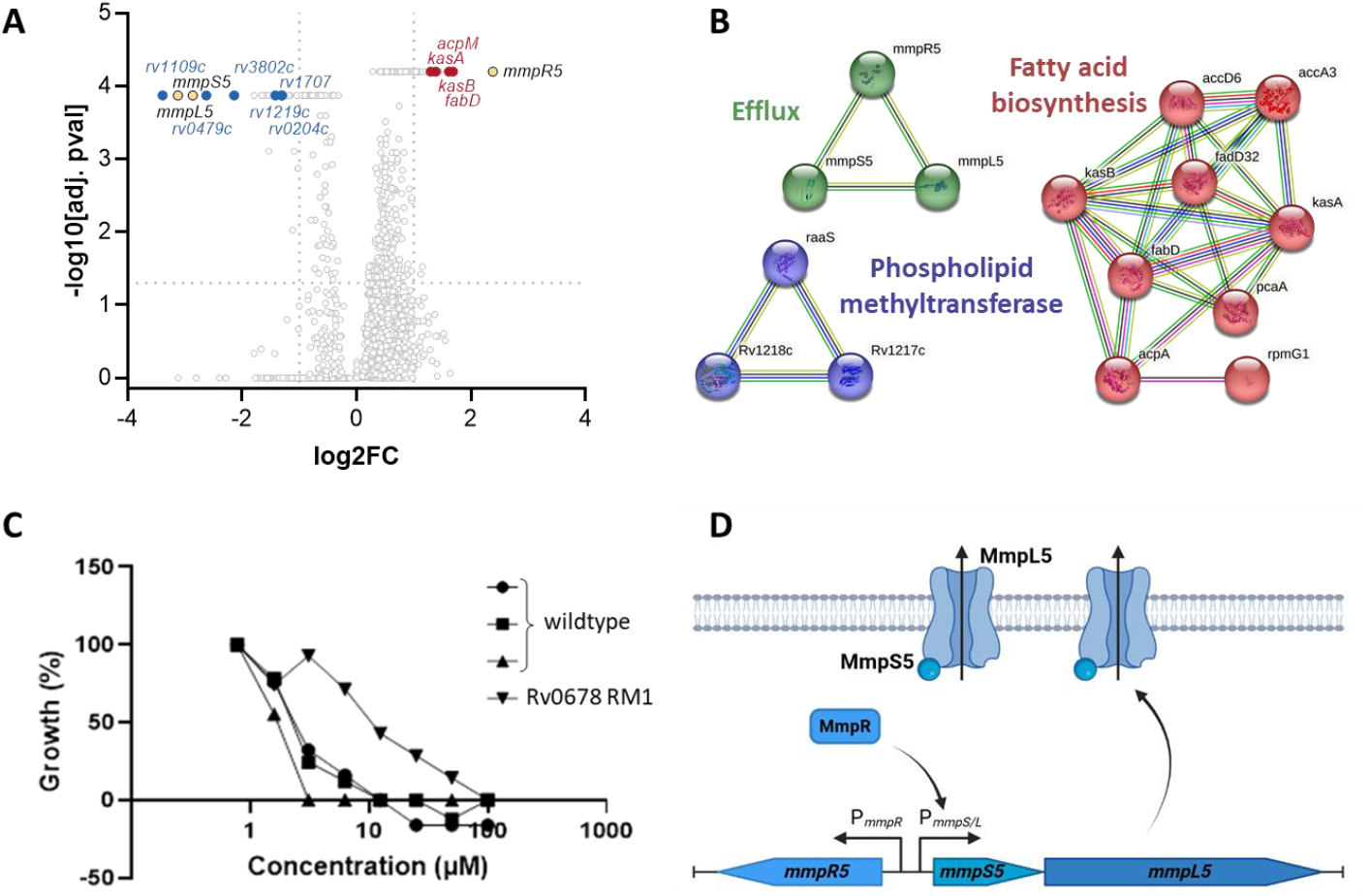
Functional genomics reveals MmpRLS5 as resistance driver for DHG. A) CRISPRi screen in M. tuberculosis shows MmpR5/MmpL5 efflux as main driver for resistance against DHG. Resistors and sensitizers are displayed to the right and left, respectively. Significant hits (p value −log10 >1.3; log2 FC>1) are colored. B) STRING network analysis of all significant CRISPRi-hits (0.5xMIC) forming distinct clusters. Individual clusters are shown in red (fatty acid biosynthesis, cluster strength 1.3), phospholipid methyl transferase (blue, cluster strength 1.4), and green (efflux system, cluster strength 1.3). Clusters were filtered based on cluster strength (>1) and false discovery rate (FDR<0.05), all non-clustering proteins have been excluded. C) Mtb H37Rv ΔRv0678 (MmpR5) is more resistant to DHG compared to its control, DREAM8, carrying wildtype MmpR5. D) Overview of the MmpL-mediated efflux system (created in BioRender. Deschner, F. (2026) https://BioRender.com/mhk5jej).

We further analyzed all significant hits from the chemical-genomic profiling (log_2_ fold-change >1) (0.5xMIC, **Table SI 2**) using the STRING database^25^ and filtered based on cluster strength and false discovery rate to gain a general overview about the assay response (**Figure 3**B). As expected, the MmpRLS5 efflux system was found with high certainty (cluster strength 1.3). We further found both the fatty acid biosynthesis and the phospholipid methyl transferase activity significantly enriched in the network (cluster strength 1.3 and 1.4, respectively). The same clusters were found when the hits from 1xMIC and 0.75xMIC DHG treatment were analyzed (**Figure SI 1**).

For hit validation, we tested Mtb H37Rv carrying *mmpR5* (Rv0678) deletions in a clean genetic background, as well as hypomorphs in two of the sensitizer genes – *rv3802c* and *rv0479c* in drug susceptibility assays. We could confirm modest susceptibility changes for all strains (**Figure 3**C/D, **Figure SI 2**): The *mmpR5* deletion mutant strain was less susceptible to DHG compared to the wildtype strain, DREAM8, confirming the relevance of the MmpRLS5-efflux system for chlorotonil resistance (**Figure 3**C/D). Conversely, hypomorphs in *rv3802c* and *rv0479c* were 2- to 5-fold more sensitive to chlorotonils compared to the non-targeting control strain (**Figure SI 2**).

### Chlorotonil-resistant Mtb H37Ra carries mutations in *mmpR5*

In an orthogonal approach, we used Mtb H37Ra to isolate mutants resistant to DHG. As previously reported, chlorotonil resistance (Ch^R^) is difficult to achieve and requires long-term exposure and adaptation (LEA)^15,26^ in liquid medium rather than direct selection for resistant clones on selective agar. Thus, we exposed Mtb H37Ra over the course of several months to DHG and slowly increased the concentration stepwise from 0.1x MIC to 1x MIC (in 50% increments). Cells treated with DMSO were used as passaging control to check for spontaneous mutations. We generated Ch^R^ Mtb with high-level resistance (32 to >64-fold shift in MIC compared to wildtype) against DHG (Table 2). After gDNA extraction, whole genome sequencing was performed, followed by genomic analysis and manual mutation calling on basis of passaging control and the H37Ra reference genome (available from ATCC genome portal).

**Table 2:** Ch^R^ M. tuberculosis shows high level resistance against DHG and carries mutations in mmpR5. Mutants were generated using a LEA-approach^26^, followed by genomic analysis. For mutants M1-M13, their MIC is displayed as fold change in relation to wildtype (WT, MIC in µg/mL is shown in brackets). Mutations are annotated as single nucleotide polymorphisms on amino acid level; ins: insertion, del: deletion; for M13, a range of nine amino acids was changed.

|  | DHG<br>MIC fold | MmpR5 | Rv2683 | Membrane<br>permease | PlsB | HddA | AdhB 2 |
| --- | --- | --- | --- | --- | --- | --- | --- |
| Mtb_WT | 1 (0.125<br>$\mu\text{g/mL}$) | | | | | | |
| Mtb_M1 | 64 |  | Met133Thr |  |  |  |  |
| Mtb_M2 | >64 | Arg156ins |  |  |  | Leu362Pro |  |
| Mtb_M3 | 32-64 | Val20Ala |  |  |  |  |  |
| Mtb_M4 | 64 | Asp8ins |  |  | Thr357Ala |  |  |
| Mtb_M5 | 64 | Ala36Thr |  |  |  |  |  |
| Mtb_M6 | 32 |  | Val143Met |  |  |  | Val258Gly |
| Mtb_M7 | 32 |  | Val75Ala |  |  |  |  |
| Mtb_M8 | 32 |  | Val75Ala |  |  |  |  |
| Mtb_M9 | 32 |  | Val75Ala |  |  |  |  |
| Mtb_M10 | >64 | Gly66del |  | Val674Ala |  |  | Ala240His<br>Glu242Stop |
| Mtb_M11 | 32 |  | Val75Ala |  |  |  |  |
| Mtb_M12 | 32-64 | Val20Ala |  |  |  |  |  |
| Mtb_M13 | 64 | Asp15-Met23<br>(multiple<br>mutations in<br>sequence) |  |  |  |  |  |

Ch^R^ Mtb primarily carried non-synonymous, deletion or insertion mutations in *mmpR5* (n=7) or non-synonymous mutations in *rv2683* (n=6) (**Table 2**) with mutations in these genes being mutually exclusive. In MmpR5, six different mutation sites were identified that spanned the entire 166 amino acid protein, whereas three distinct mutations were found in Rv2683. The mutations in *mmpR5* generally conferred higher level resistance (32 to >64-fold) than those in *rv2683* (32-fold). While seven out of 13 analyzed mutants confirmed MmpR5 as driver for Ch^R^, the role of the conserved, membrane-associated protein Rv2683 found in the other six mutants remains elusive, as direct enzymatic or transport roles remain undefined; studying the role and function of Rv2683 was beyond the scope of this study but will be addressed in future experiments.

We further identified mutations in an uncharacterized membrane permease, *plsB* (Glycerol-3-phosphate acyltransferase), *hddA* (D-glycero-alpha-D-manno-heptose 7-phosphate kinase) and *adhB 2* (alcohol dehydrogenase). However, these alterations showed no clear association with resistance phenotypes, as 9 of the 13 mutants already exhibited high-level resistance solely through mutations in MmpR5 or Rv2683, raising questions about the functional contribution of these less frequently observed mutations, hence, we did not further elaborate (**Table 2**).

### MmpR5-mediated chlorotonil-resistance also confers resistance to BDQ

As *mmpR5* mutations are known to confer resistance to last-line antibiotic BDQ^27^, we analyzed our Ch^R^-mutants carrying *mmpR5* mutations for cross-resistance against BDQ and the reference drugs rifampicin (RIF) and isoniazid (INH) (**Table 3**). We confirmed cross-resistance of Ch-induced resistance also towards BDQ, while no change in susceptibility could be observed for RIF and INH. Interestingly, the exchange of Val with Ala at position 20 (Val20Ala), resulted only in a 4-fold reduced susceptibility to BDQ, compared to 8-fold for the other mutations.

**Table 3:** Chlorotonil-resistant M. tuberculosis is also resistant to BDQ. Regular Mtb H37Ra and passaging control (Mtb WT mutants) were used as reference for Ch^R^-mutants M2-5, M10, M12 and M13 carrying mutations in mmpR5; respective mutations in mmpR5 can be seen in each column.

| Compound | MIC [μg/mL] |  |  |  |  |  |  |  |  |
| --- | --- | --- | --- | --- | --- | --- | --- | --- | --- |
|  | Mtb H37Ra | Mtb WT mutants | M2 R156ins | M3 V20A | M4 D8ins | M5 A36T | M10 G66del | M12 V20A | M13 A15-M23 |
| DHG | 0.25 | 0.25 | > 8 | 4 | > 8 | > 8 | > 8 | 4 | > 8 |
| BDQ | 0.0156 | 0.0313 | 0.25 | 0.125 | 0.25 | 0.25 | 0.25 | 0.125 | 0.5 |
| RIF | 0.0156 | 0.0313 | 0.0313 | 0.0313 | 0.0313 | 0.0313 | 0.0313 | 0.0313 | 0.0313 |
| INH | 0.5 | 0.5 | 0.25 | 0.5 | 0.25 | 0.5 | 0.5 | 0.5 | 0.5 |

To further investigate a potential mechanistic similarity of chlorotonils to BDQ, we compared the DHG ad BDQ susceptibilities of *atpB* hypomorphs and an AtpE^Ala63Pro^ mutant strain of Mtb, which serve as reporters for on-target ATP synthase inhibitors. While the strains showed the expected susceptibility shifts to BDQ – hypersensitivity of the *atpB* hypomorphs vs. increased resistance of the AtpE^Ala63Pro^ mutant – they showed no differential susceptibility to DHG, this arguing against ATP synthase as a direct target of DHG (**Figure SI 3**).

## 3. Discussion

Natural product chlorotonils have been shown to be effective against several Gram-positive bacteria, and in this study, we characterized their activity against *Mycobacterium* spp. The MIC of chlorotonils against Mtb is similar to that observed for Gram-positives^13,14^, suggesting that these molecules can traverse the lipid-rich cell envelope to reach their cellular target/s in Mtb – typically a challenge in TB drug discovery^28,29^. In contrast, chlorotonils are inactive against NTMs. Time-kill kinetics revealed a relatively slow rate of kill of DHG against Mtb at a high drug concentration unlike the ultrafast kinetic profile reported for *S. aureus, Clostridioides difficile* or *P. falciparum*^15,16,30^. Although Myco-BCP analysis found a high similarity score to the PMF inhibitor, CCCP, and the ATP synthase inhibitor, BDQ, we could not confirm direct chlorotonil-mediated membrane depolarization in Mtb, which was identified as primary reason for the anti-staphylococcal activity of DHG^15^. While CCCP is a classical ionophore that directly collapses membrane potential, BDQ inhibits ATP synthase and depletes ATP without immediate membrane depolarization^19,20^. Generally, Mtb appears to be less sensitive to membrane depolarizing effects, which may be attributed to its slow metabolism as slow-growing bacteria tend to be more tolerant to such antibiotic stressors^31–34^. Thus, even if chlorotonils reach the membrane but do not cause measurable depolarization, they could still exert an antimycobacterial effect through other pathways, *e.g.*, perturbation of membrane-associated processes such as transport systems or cellular respiration and energy-metabolism. The similarity in cytological profiles therefore reflects convergence on impaired energy production – similar to BDQ – and the profiling data suggest chlorotonils are most consistent with this pathway.

The interplay between energy metabolism and lipid metabolism in Mtb is critical for its survival, adaptation, and response to drug treatment. Lipid metabolism can be considered an “energy-sink” due to how Mtb stores and mobilizes lipids for energy, which directly affects its drug sensitivity^35,36^. In turn, slowing down lipid metabolism and ATP consumption would confer resistance when energy-metabolism is targeted. Two lines of evidence implicated lipid metabolism in the antimycobacterial MoA of chlorotonils: (i) DHG treatment of *M. smegmatis* elicited a morphotype most closely related to that observed for hypomorphs in lipid metabolism; (ii) the partial knockdown of the lipid biosynthesis genes, *fas, accA3, acpM, kasA/B, fabD*, reduced the susceptibility of Mtb to chlorotonil whereas partial knockdown of *rv3802c*, which encodes a phospholipase/thioesterase implicated in the MoA of tetrahydrolipstatin and is linked to mycolic acid biosynthesis and glycerophospholipid composition in the membrane^37,38^ resulted in modest sensitization to DHG. In Ch^R^ mutants we also found mutations in Rv2683, PlsB, HddA, and AdhB2, and while their molecular role in Ch^R^ remains elusive, all proteins can be connected to lipid metabolism. They play a role during cholesterol uptake and glycerophospholipid metabolism and are, for example, directly responsible for the synthesis of cell envelope components^39–46^.

At this stage we cannot fully describe the mechanism of chlorotonils in Mtb, but we have strong evidence that both lipid and energy metabolism play a role, which fits the lipid-mediated activity in other species. Likewise, the mechanism of DHG in Mtb appears complex but distinct from that in *S. aureus* where, *e.g*., two additional enzyme targets involved in cell wall and protein biosynthesis have been identified^15^. Activity against NTM species needs to be investigated in the future, as we currently lack a clear explanation for the complete inactivity of chlorotonils. Several factors could contribute to this phenotype, including differences in cell wall lipid composition that reduce permeability or result in distinct lipid structures, as well the high abundance of efflux pumps and intrinsic resistance mechanisms in these mycobacterial species^47–49^.

We identified the MmpS5-MmpL5 efflux system as a key contributor to chlorotonil resistance in Mtb using a combination of resistance mutant generation and chemical-genomic profiling. Mutations in the repressor protein MmpR5 with subsequent upregulation of MmpL5-efflux are also known to confer resistance to BDQ and other important anti-TB drugs including clofazimine and benzothiazinone DprE1 inhibitors^8,27,50,51^. In an otherwise isogenic background, deletion of *mmpR5* resulted in an approximately five-fold reduced susceptibility of Mtb to DHG compared to the wildtype strain, consistent with previously reported resistance levels (4-to 8-fold) associated with *mmpR5* mutations in BDQ^R^ Mtb strains^8,27^. Strikingly, Ch^R^ Mtb isolates exhibited more than 64-fold reduced susceptibility, indicating that additional mechanisms beyond MmpL-mediated efflux may contribute to high-level resistance. At this stage, we cannot rule out further contributions by Rv2683 or other observed mutations that are for example linked to lipid metabolism. Notably, in *S. aureus* FarE was found to be responsible for high-level resistance against chlorotonil in a similar way as reported for rhodomyrtone^15,52^. FarE pumps lipids and fatty acids across the membrane, which then in turn capture and neutralize DHG molecules resulting in compound “detoxification” outside the cell. FarE belongs to the same major MmpL-protein family as the MmpS5-MmpL5 pump, hence there is the possibility that resistance to chlorotonils in Mtb is also caused by fatty acid/lipid export and capture resulting in “detoxification” rather than being solely caused by drug-efflux via MmpL, as reported for BDQ^8^. We speculate that in NTM species, a similar mechanism may cause resistance of the wild-type strains. Obviously, more research is necessary to understand MmpL5 and its substrates to evaluate such theory in the future.

The confirmation of co-resistance to BDQ as one of the corner drugs for MDR-TB therapy led us to deprioritize chlorotonils as candidates for TB drug discovery and development. However, these compounds have the potential for use as tool compounds for investigating the biological function (and possible opportunities for disruption) of the MmpS5-MmpL5 pump given its recognized role in compromising the clinical efficacy of BDQ.

## 4. Acknowledgements

The authors want to thank Arwa Alharbi and Xiaoxin An for help with processing samples for CRISPRi profiling and Timothy de Wet for analyzing morphological profiling data in *M. smegmatis*. This work was supported by the Gates Foundation (INV-055896 to DS, INV-055894 to DS and JR, INV-040484 to VM, INV-055900 to RM).

## 5. Materials and Methods

### Cultivation and susceptibility testing

Mtb H37Ra (ATCC25177) was retrieved from the American Type Culture Collection (ATCC) and grown at 37°C in Middelbrook 7H9 medium supplemented according to manufactures instructions. Mtb H37Rv was used for all CRISPRi experiments and grown at 37°C in Difco Middlebrook 7H9 broth (BD #271310) or 7H10 (BD #262710) plates supplemented with 0.2% glycerol (7H9) or 0.5% glycerol (7H10), 0.05% Tween80 and 10% Oleic-Albumin-Dextrose-Catalase (OADC). When required, kanamycin was used at 10-20 µg/mL and ATc at 100 ng/mL.

For susceptibility testing, cells were grown to early logarithmic phase (OD0.4-0.8), followed by single cell preparation using needle extrusion according to internal standard protocols. Testing was performed in M7H9 medium without the addition of Tween20, with a starting inoculum of OD0.001 (1.25×10^5^ CFU/mL). Compounds were tested in 10 different concentrations (1/2 dilutions) starting at 8 µg/mL (DHG, ChB1-Epo2) or 1.6 µg/mL (ChA). Plates were incubated at 37°C for 7 days and evaluated using OD or resazurin (final concentration 0.002%).

For NTM (including MAB smooth and rough and MAH), Resazurin microtiter assay (REMA) was performed in 96-well plates with *M. abscessus* subsp. *abscessus* ATCC 19977 smooth and rough morphotypes and *M. avium* subsp. *hominissuis* strain 104^53^. First, a mid-logarithmic-phase culture (OD approximately 0.5) was diluted in 7H9 complete medium with Tween80 to an OD600 of 0.001 (approximately 5×10^5^ CFU/mL). Bacteria (100 μL) were then dispensed in transparent flat-bottom 96-well plates. Two-fold serial dilutions of each drug were then prepared. On each plate, controls without drug and medium only were included. The plates were incubated for 2 days at 37°C before adding the resazurin (0.025% [*w/v*] to 1/10 of the well volume). After overnight incubation, the fluorescence of the resazurin metabolite, resorufin, was determined with excitation at 560 nm and emission at 590 nm, measured using a Tecan Infinite M Plex microplate reader. The MIC was calculated using GraphPad Prism software (version 9) and the Gompertz equation for MIC determination. All drugs were tested at least in biological triplicates.

### Time-kill curve

Bacteria were prepared as for regular MIC testing and exposed to 1 µg/mL or 0.1 µg/mL DHG (or DMSO as control). At indicated time points, 100 µL were taken and diluted in 50% FBS/MQ and plated on standard Middlebrook 7H10 agar. After 4-6 weeks incubation in sealed bags at 37°C, colonies were counted.

### Membrane depolarization

Bacterial cells were grown in M7H9+OADC+Tween20 until OD0.4, then 0.1% glucose was added to the medium and grown until OD0.6-0.8^15^. Cells were collected, supernatant removed and resuspended in PBS+0.1% glucose. Cells were stained for 30 min in the dark using 30 µM DiOC_2_(3) and 100 µL were transferred per well into a black 96 flat-bottom well plate, prior measurement of the plate using a Tecan Spark (green: 485/530 nm; red 485/630 nm). After 4 min of baseline recording, test compounds (including DMSO) were added to a final DMSO concentration of 1% (n=2) and reading was continued until 30 min. For evaluation, the red/green ratio was normalized against the respective DMSO time points.

### BCP

MycoBCP was performed as described by Quach et al^54^. To prepare cultures for BCP, we diluted a parent *M. tuberculosis* culture grown in 7H9+PL at 30 °C to a final OD_600_ of ∼0.06 to 0.08 which was then rolled at 30 °C for 18 to 20 h, a slower growth temperature that provided reproducible phenotypes. After this outgrowth, the culture was split into 2 mL aliquots and treated with sufficient compound to reach the desired test concentration, typically between 1× and 5× the MIC. These treated aliquots were then rolled at 30 °C, and 400 μL samples were harvested and fixed after 48 and 120 h of treatment. Specifically, each 400 μL sample was fixed at 30 °C for 20 min with a mixture of 100 μL of 16% paraformaldehyde, 3 μL of 8% glutaraldehyde, and 20 μL of 0.4 M phosphate buffer, pH 7.5. After fixation, the cells were washed twice with 200 μL of warm 7H9+PL concentrated to a final volume of ∼30 μL and stained for 30 min. The cells were stained with 0.04 mg/mL FM 4-64 (Invitrogen), 0.1 to 0.2 mM SYTO 40 (Invitrogen), and 4 µM SYTOX Green (Invitrogen). In images, membranes are shown in red, DNA in blue, and green staining indicates that cell integrity has been compromised; white scale bars are 1 µm. For image preparation, full-field fluorescence microscopy images of samples prepared for BCP are preprocessed from their original proprietary microscope file format to Tagged Image File Format. The original image dimensions 3 × 2,048 × 2,048 are cropped to 3 × 1,800 × 1,800 by trimming 124 pixels from each edge to avoid optical artifacts. Each image is divided into nine subimages of dimensions 3 × 600 × 600. Image entropy [calculated as sum(p*log2(p)] where p is the normalized histogram counts, is used to automatically filter out images without a significant number of cells.

### Morphological profiling of *M. smegmatis*

Sample preparation and imaging were performed as described by de Wet et al. (2020)^21^. Briefly, a *M. smegmatis* parB-mCherry fluorescent labeled strain (chromosome partitioning marker strain) was treated with compound for 18 h at 1, 2 or 4X MIC. A no-drug control was included for comparison. Following treatment, the bacilli were spotted onto agarose pads and images acquired using a Zeiss Axio Observer Z1 microscope and ZEN 2 (blue edition) software. Approximately 20 random fields of view were captured with a 100X Phase Contrast Objective (1.4NA). Images were processed in FIJI^55^ and analyzed using custom scripts in Microbe J^56^. Thirteen morphological features were recorded and converted into a Z-score which was plotted onto a Uniform manifold approximation and projection (UMAP) map containing clusters of orthologous groups (COG) annotations.

### ATP depletion assay

*M. tuberculosis* H37Rv-LP was grown to late log phase and inoculated at OD_590_0.04 in 96-well plates containing compounds as two-fold dilutions. Plates were incubated at 37°C for 24 hours. ATP was quantified by adding 50 µL BacTiter-Glo™ reagent per well, incubating for 10 min and reading luminescence using a Synergy H4 plate reader. Kanamycin and Q203 were used as negative and positive controls, respectively.

### CRISPRi library design, construction and profiling

The CRISPRi Library (RLC25; Addgen) was designed to target all possible *M. tuberculosis* ORFs and non-coding RNAs. RLC25 is a combination of two libraries: 1) RLC3, which was previously designed to target predicted *in vitro* non-essential genes^57^, and 2) RLC14, which is a library designed to target in-vitro essential genes with hypomorphic sgRNAs of different strengths.

To design hypomorphic sgRNAs for library RLC14, we started with estimates of the rate of depletion of sgRNAs according to a piecewise (“two-line”) model as described in Bosch & DeJesus et al. For each *in vitro* essential, a maximum of 20 sgRNAs was selected. sgRNAs were selected to prioritize depletion rates around Beta_E = −0.14, while including some stronger and weaker guides. To achieve this, bins representing the 20^th^ percentiles of a normal distribution centered at –0.14 were defined, and then a single guide from each bin was selected. This helped guarantee that most sgRNA’s would be selected around –0.14 (where the mass of the distribution, and thus percentiles, lies) while including some strong and weak guides at the top and bottom percentiles. If a bin was lacking sgRNAs, other bins would be repeated when possible until a maximum of 20 sgRNAs was achieved. A total of 14,354 guides targeting 748 essential genes were selected using this approach, with 646 non-targeting guides added for a total of 15,000 guides in RLC14. After combining RLC3 and RLC14, the final library contained a total of 30,306 guides targeting a total of 4,005 genes and including 837 non-targeting control sgRNAs.

For each profiling experiment, 1 mL aliquots of RLC0025 (OD_580_ ∼1 per mL) were used to inoculate 24 mL 7H9 supplemented with kanamycin (10 μg/mL) in vented 75 cm^2^ tissue culture flasks. Cultures were expanded for 5 days to OD_580_=∼1.2, pooled and passed through a 10 μm cell strainer to obtain a single-cell suspension. The single-cell suspension was diluted to OD_580_=0.5 in a total volume of 75 mL 7H9 with ATc (100 ng/mL) and incubated at 37°C for another 24 h. Triplicate cultures were then inoculated at OD_580_=0.05 in 3.3 mL 7H9 supplemented with ATc, kanamycin (10 μg/mL), and the compound under investigation (dissolved in DMSO) or DMSO without compound. Cultures were grown in vented 12.5 cm^2^ tissue culture flasks for 14 days at 37°C, 5% CO2. ATc was replenished at day 7. After 14 days of incubation, OD_580_ values were measured and cultures were harvested. Genomic DNA was isolated from bacterial pellets using the Omega Bio-Tek Mag-Bind Universal Pathogen DNA Kit (M4029), along with a Geno/Grinder for cell lysis and a Hamilton Starlet for partial automation of the extraction process. The concentration of isolated genomic DNA was quantified using a Thermo Scientific NanoDrop ND-8000 Spectrophotometer. Subsequently, the sgRNA-encoding region was amplified, purified and pooled for Illumina sequencing as previously described in Li et al. (2022).^23^

### Hypomorphs

AtpB, Rv0479c and Rv3802 hypomorphs were created as previously described in Chengalroyen et al. (2024) utilizing the anhydrotetracycline (ATc)-regulated CRISPRi system developed by Rock and colleagues^58,59^. Briefly, the hypomorphs and non-target (NT) sgRNA control strains were grown to an OD600 of 1.0 and diluted 10X in media supplemented with 100 ng/ml ATc. This was incubated for 3 days (ATc pre-depletion step). All cultures were adjusted to an OD600 of 1.0, diluted 20X and 50 μl of the diluted culture inoculated into each well of a MIC plate containing 50 μl of media with 2-fold dilutions of compound and 100 ng/ml ATc. The microtiter plates were sealed and incubated at 37°C for 14 days and the ODs were recorded using the SpectraMax i3x plate reader. Each strain was normalized to the no-drug control. Accordingly, the percentage growth inhibition was plotted as a function of drug concentration using Prism 9 (GraphPad).

### Resistance generation

Resistance was generated by adaptation similar to previously described methods^26^. In brief, twenty 5 mL cultures were exposed to sub-inhibitory concentrations of DHG (start: 0.1xMIC) in M7H9+OADC without tween and grown until sufficient turbidity was reached. Then, cultures were split 1/33 into fresh medium containing 50% increased concentration of DHG. A passaging control exposed to DMSO only was kept and treated equally to control for random mutation events.

### Extraction and analysis of genomic DNA

Genomic DNA was extracted using the CTAB-Chloroform method. 50 mL culture was centrifuged and resuspended in 1 mL TE buffer (10 mM Tris/HCl pH 8, 1 mM EDTA, 25 mM NaCl). Cell lysis was performed overnight at 37°C using 100 µL lysozyme (20 mg/mL). Then, 2.75 µL RNAse (10 mg/mL) was added and incubated for 1h (37°C) followed by addition of 50 µL proteinase K (10 mg/mL) and incubation at 65°C for 2h (shaking). 1 mL of suspension was transferred to a fresh tube, mixed with 180 µL preheated (65°C) 5M NaCl and 100 µL CTAB (10%), and immediately thoroughly mixed. Samples were kept at 65°C for 15 min (shaking) and then frozen at −80°C for 15 min, followed by thawing at room temperature for 10 min and further incubation at 65°C for 15 min. Tubes were cooled to room temperature prior addition of 700 µL chloroform:isoamyl alcohol (24:1). Tubes were kept for 10 min on inverter (40 rpm), then left for 10 min on the bench to allow for proper phase separation, followed by 10 min centrifugation at 13,000 rpm. Upper aqueous phase was then transferred to fresh tube containing 1 volume of fresh ice-cold isopropanol and thoroughly mixed. Tubes were kept at −20°C overnight and then centrifuged at 4°C to precipitate DNA. Samples were drained and pellet was washed using 800 µL cold 80% ethanol. After centrifugation, pellet was dried in speed vac (low temperature) for 10 min and then dissolved in 55 µL TE buffer prior to storage at −20°C.

The gDNA of wildtype and mutant samples were sent for Illumina MiSeq (2×300 bp reads, coverage >100) and analyzed using Geneious Prime Version 2022. In brief, paired reads were imported and mapped against the respective annotated reference genome retrieved from ATCC Genome Portal for *M. tuberculosis* ATCC25177. After mapping, the consensus sequences were generated and aligned against the annotated reference genome (Progressive Mauve Algorithm) with a match seed weight of 15 and a minimum LCB score of 30,000. SNP calling was performed manually on the bases of both the original wildtype sample and the sequencing control grown for the entirety of the adaptation process.

WGS data are available at the National Center for Biotechnology Information (NCBI) through the BioProject accession number PRJNA1354405.

## Notes

### Competing Interest Statement

The authors have declared no competing interest.

## Bibliography

1. World Health Organization (2025). Global tuberculosis report 2025.

2. Warner, D.F., Barczak, A.K., Gutierrez, M.G., and Mizrahi, V. (2025). Mycobacterium tuberculosis biology, pathogenicity and interaction with the host. Nat Rev Microbiol 23, 788– 804. 10.1038/s41579-025-01201-x.

3. Yu, X., Liu, P., Liu, G., Zhao, L., Hu, Y., Wei, G., Luo, J., and Huang, H. (2016). The prevalence of non-tuberculous mycobacterial infections in mainland China: Systematic review and meta-analysis. J Infect 73, 558–567. 10.1016/J.JINF.2016.08.020.

4. Duarte, R., Munsiff, S.S., Nahid, P., Saukkonen, J.J., Winston, C.A., Abubakar, I., Acuña-Villaorduña, C., Barry, P.M., Bastos, M.L., Carr, W., et al. (2025). Updates on the Treatment of Drug-Susceptible and Drug-Resistant Tuberculosis An Official ATS/CDC/ERS/IDSA Clinical Practice Guideline. Am J Respir Crit Care Med 211, 15–33. 10.1164/RCCM.202410-2096ST.

5. World Health Organization (2022). WHO consolidated guidelines on tuberculosis. Module 4: treatment - drug-resistant tuberculosis treatment, 2022 update.

6. Motta, I., Boeree, M., Chesov, D., Dheda, K., Günther, G., Horsburgh, C.R., Kherabi, Y., Lange, C., Lienhardt, C., McIlleron, H.M., et al. (2024). Recent advances in the treatment of tuberculosis. Clin Microbiol Infect 30, 1107–1114. 10.1016/j.cmi.2023.07.013.

7. Chahine, E.B., Karaoui, L.R., and Mansour, H. (2014). Bedaquiline: A Novel Diarylquinoline for Multidrug-Resistant Tuberculosis. Annals of Pharmacotherapy 48, 107–115. 10.1177/1060028013504087.

8. Andries, K., Villellas, C., Coeck, N., Thys, K., Gevers, T., Vranckx, L., Lounis, N., De Jong, B.C., and Koul, A. (2014). Acquired resistance of Mycobacterium tuberculosis to bedaquiline. PLoS One 9. 10.1371/JOURNAL.PONE.0102135.

9. World Health Organization (2024). WHO bacterial priority pathogens list, 2024: bacterial pathogens of public health importance, to guide research, development and strategies to prevent and control antimicrobial resistance (World Health Organization).

10. WHO (2017). WHO | Global priority list of antibiotic-resistant bacteria to guide research, discovery, and development of new antibiotics. WHO, 1–7.

11. Newman, D.J., and Cragg, G.M. (2020). Natural Products as Sources of New Drugs over the Nearly Four Decades from 01/1981 to 09/2019. J Nat Prod 83, 770–803.

12. Held, J., Gebru, T., Kalesse, M., Jansen, R., Gerth, K., Müller, R., and Mordmüller, B. (2014). Antimalarial Activity of the Myxobacterial Macrolide Chlorotonil A. Antimicrob Agents Chemother 58, 6378. 10.1128/AAC.03326-14.

13. Hofer, W., Oueis, E., Fayad, A.A., Deschner, F., Andreas, A., de Carvalho, L.P., Hüttel, S., Bernecker, S., Pätzold, L., Morgenstern, B., et al. (2022). Regio- and Stereoselective Epoxidation and Acidic Epoxide Opening of Antibacterial and Antiplasmodial Chlorotonils Yield Highly Potent Derivatives. Angewandte Chemie International Edition 61, e202202816. 10.1002/ANIE.202202816.

14. Hofer, W., Deschner, F., Jézéquel, G., Pessanha de Carvalho, L., Abdel-Wadood, N., Pätzold, L., Bernecker, S., Morgenstern, B., Kany, A.M., Große, M., et al. (2024). Functionalization of Chlorotonils: Dehalogenil as Promising Lead Compound for In Vivo Application. Angewandte Chemie International Edition 63, e202319765. 10.1002/ANIE.202319765.

15. Deschner, F., Mostert, D., Daniel, J.-M., Voltz, A., Schneider, D.C., Khangholi, N., Bartel, J., Pessanha de Carvalho, L., Brauer, M., Gorelik, T.E., et al. (2025). Natural products chlorotonils exert a complex antibacterial mechanism and address multiple targets. Cell Chem Biol 32, 586-602.e15. 10.1016/j.chembiol.2025.03.005.

16. de Carvalho, L.P., Niepoth, E., Mavraj-Husejni, A., Kreidenweiss, A., Herrmann, J., Müller, R., Knaab, T., Burckhardt, B.B., Kurz, T., and Held, J. (2023). Quantification of Plasmodium falciparum HRP-2 as an alternative method to [3H]hypoxanthine incorporation to measure the parasite reduction ratio in vitro. Int J Antimicrob Agents 62, 106894. 10.1016/J.IJANTIMICAG.2023.106894.

17. Deschner, F., Mostert, D., Daniel, J.-M., Voltz, A., Schneider, D.C., Khangholi, N., Bartel, J., Pessanha de Carvalho, L., Brauer, M., Gorelik, T.E., et al. (2025). Natural products chlorotonils exert a complex antibacterial mechanism and address multiple targets. Cell Chem Biol 32, 586-602.e15. 10.1016/j.chembiol.2025.03.005.

18. Nonejuie, P., Burkart, M., Pogliano, K., and Pogliano, J. (2013). Bacterial cytological profiling rapidly identifies the cellular pathways targeted by antibacterial molecules. Proceedings of the National Academy of Sciences 110, 16169–16174. 10.1073/pnas.1311066110.

19. Lamprecht, D.A., Finin, P.M., Rahman, M.A., Cumming, B.M., Russell, S.L., Jonnala, S.R., Adamson, J.H., and Steyn, A.J.C. (2016). Turning the respiratory flexibility of Mycobacterium tuberculosis against itself. Nat Commun 7, 12393. 10.1038/NCOMMS12393.

20. Harrison, S.H., Walters, R.C., Cheung, C.-Y., Springett, R.J., Cook, G.M., Osman, M.M., and Blaza, J.N. (2025). Remission spectroscopy resolves the mechanism of action of bedaquiline within living mycobacteria. Nature Communications 2025. 10.1038/S41467-025-65928-0.

21. de Wet, T.J., Winkler, K.R., Mhlanga, M., Mizrahi, V., and Warner, D.F. (2020). Arrayed crispri and quantitative imaging describe the morphotypic landscape of essential mycobacterial genes. Elife 9, 1–36. 10.7554/ELIFE.60083.

22. Bahuguna, A., Rawat, S., and Rawat, D.S. (2021). QcrB in Mycobacterium tuberculosis: The new drug target of antitubercular agents. Med Res Rev 41, 2565–2581. 10.1002/MED.21779.

23. Li, S., Poulton, N.C., Chang, J.S., Azadian, Z.A., DeJesus, M.A., Ruecker, N., Zimmerman, M.D., Eckartt, K.A., Bosch, B., Engelhart, C.A., et al. (2022). CRISPRi chemical genetics and comparative genomics identify genes mediating drug potency in Mycobacterium tuberculosis. Nature Microbiology 2022 7:6 7, 766–779. 10.1038/s41564-022-01130-y.

24. Rodriguez, R., Campbell-Kruger, N., Camba, J.G., Berude, J., Fetterman, R., and Stanley, S. (2023). MarR-Dependent Transcriptional Regulation of mmpSL5 Induces Ethionamide Resistance in Mycobacterium abscessus. Antimicrob Agents Chemother 67. 10.1128/AAC.01350-22.

25. Szklarczyk, D., Kirsch, R., Koutrouli, M., Nastou, K., Mehryary, F., Hachilif, R., Gable, A.L., Fang, T., Doncheva, N.T., Pyysalo, S., et al. (2023). The STRING database in 2023: protein–protein association networks and functional enrichment analyses for any sequenced genome of interest. Nucleic Acids Res 51, D638–D646. 10.1093/nar/gkac1000.

26. Deschner, F., Risch, T., Baier, C., Schlüter, D., Herrmann, J., and Müller, R. (2023). Nitroxoline resistance is associated with significant fitness loss and diminishes in vivo virulence of Escherichia coli. Microbiol Spectr 12, e0307923. 10.1128/spectrum.03079-23.

27. Roberts, L.W., Malone, K.M., Hunt, M., Joseph, L., Wintringer, P., Knaggs, J., Crook, D., Farhat, M.R., Iqbal, Z., and Omar, S. V (2024). MmpR5 protein truncation and bedaquiline resistance in Mycobacterium tuberculosis isolates from South Africa: a genomic analysis. Lancet Microbe. 10.1016/S2666-5247(24)00053-3.

28. Jacobo-Delgado, Y.M., Rodríguez-Carlos, A., Serrano, C.J., and Rivas-Santiago, B. (2023). Mycobacterium tuberculosis cell-wall and antimicrobial peptides: a mission impossible? Front Immunol 14. 10.3389/FIMMU.2023.1194923.

29. Belete, T.M. (2022). Recent Progress in the Development of Novel Mycobacterium Cell Wall Inhibitor to Combat Drug-Resistant Tuberculosis. Microbiol Insights 15, 117863612210998. 10.1177/11786361221099878.

30. Bublitz, A., Brauer, M., Wagner, S., Hofer, W., Müsken, M., Deschner, F., Lesker, T.R., Neumann-Schaal, M., Paul, L.S., Nübel, U., et al. (2023). The natural product chlorotonil A preserves colonization resistance and prevents relapsing Clostridioides difficile infection. Cell Host Microbe 31, 734-750.e8. 10.1016/J.CHOM.2023.04.003.

31. Hennes, M., Bender, N., Cronenberg, T., Welker, A., and Maier, B. (2023). Collective polarization dynamics in bacterial colonies signify the occurrence of distinct subpopulations. PLoS Biol 21. 10.1371/JOURNAL.PBIO.3001960.

32. Grassi, L., Di Luca, M., Maisetta, G., Rinaldi, A.C., Esin, S., Trampuz, A., and Batoni, G. (2017). Generation of Persister Cells of Pseudomonas aeruginosa and Staphylococcus aureus by Chemical Treatment and Evaluation of Their Susceptibility to Membrane-Targeting Agents. Front Microbiol 8, 1917. 10.3389/fmicb.2017.01917.

33. Benarroch, J.M., and Asally, M. (2020). The Microbiologist’s Guide to Membrane Potential Dynamics. Preprint at Elsevier Ltd, 10.1016/j.tim.2019.12.008 10.1016/j.tim.2019.12.008.

34. Stokes, J.M., Lopatkin, A.J., Lobritz, M.A., and Collins, J.J. (2019). Bacterial Metabolism and Antibiotic Efficacy. Cell Metab 30, 251. 10.1016/J.CMET.2019.06.009.

35. Kim, H., and Shin, S.J. (2023). Revolutionizing control strategies against Mycobacterium tuberculosis infection through selected targeting of lipid metabolism. Cell Mol Life Sci 80, 291. 10.1007/s00018-023-04914-5.

36. Rodríguez, J.G., Hernández, A.C., Helguera-Repetto, C., Aguilar Ayala, D., Guadarrama-Medina, R., Anzóla, J.M., Bustos, J.R., Zambrano, M.M., González-y-Merchand, J., García, M.J., et al. (2014). Global Adaptation to a Lipid Environment Triggers the Dormancy-Related Phenotype of Mycobacterium tuberculosis. mBio 5. 10.1128/mBio.01125-14.

37. Crellin, P.K., Vivian, J.P., Scoble, J., Chow, F.M., West, N.P., Brammananth, R., Proellocks, N.I., Shahine, A., Le Nours, J., Wilce, M.C.J., et al. (2010). Tetrahydrolipstatin Inhibition, Functional Analyses, and Three-dimensional Structure of a Lipase Essential for Mycobacterial Viability. Journal of Biological Chemistry 285, 30050–30060. 10.1074/jbc.M110.150094.

38. Parker, S.K., Barkley, R.M., Rino, J.G., and Vasil, M.L. (2009). Mycobacterium tuberculosis Rv3802c Encodes a Phospholipase/Thioesterase and Is Inhibited by the Antimycobacterial Agent Tetrahydrolipstatin. PLoS One 4, e4281. 10.1371/journal.pone.0004281.

39. Mir, M., Prisic, S., Kang, C.M., Lun, S., Guo, H., Murry, J.P., Rubin, E.J., and Husson, R.N. (2014). Mycobacterial Gene cuvA Is Required for Optimal Nutrient Utilization and Virulence. Infect Immun 82, 4104. 10.1128/IAI.02207-14.

40. Brzostek, A., Pawelczyk, J., Rumijowska-Galewicz, A., Dziadek, B., and Dziadek, J. (2009). Mycobacterium tuberculosis is able to accumulate and utilize cholesterol. J Bacteriol 191, 6584–6591. 10.1128/JB.00488-09.

41. Chen, Z., Kong, X., Ma, Q., Chen, J., Zeng, Y., Liu, H., Wang, X., and Lu, S. (2024). The impact of Mycobacterium tuberculosis on the macrophage cholesterol metabolism pathway. Front Immunol 15. 10.3389/FIMMU.2024.1402024.

42. Karan, S., Behl, A., Sagar, A., Bandyopadhyay, A., and Saxena, A.K. (2021). Structural studies on Mycobacterium tuberculosis HddA enzyme using small angle X-ray scattering and dynamics simulation techniques. Int J Biol Macromol 171, 28–36. 10.1016/J.IJBIOMAC.2020.12.191.

43. Bourai, N., Jacobs, W.R., and Narayanan, S. (2012). Deletion and overexpression studies on DacB2, a putative low molecular mass penicillin binding protein from Mycobacterium tuberculosis H(37)Rv. Microb Pathog 52, 109–116. 10.1016/J.MICPATH.2011.11.003.

44. Nazarova, E. V., Montague, C.R., La, T., Wilburn, K.M., Sukumar, N., Lee, W., Caldwell, S., Russell, D.G., and VanderVen, B.C. (2017). Rv3723/LucA coordinates fatty acid and cholesterol uptake in Mycobacterium tuberculosis. Elife 6. 10.7554/ELIFE.26969.

45. Griffin, J.E., Gawronski, J.D., DeJesus, M.A., Ioerger, T.R., Akerley, B.J., and Sassetti, C.M. (2011). High-Resolution Phenotypic Profiling Defines Genes Essential for Mycobacterial Growth and Cholesterol Catabolism. PLoS Pathog 7, e1002251. 10.1371/journal.ppat.1002251.

46. Chen, Y., Hagopian, B., and Tan, S. (2025). Cholesterol metabolism and intrabacterial potassium homeostasis are intrinsically related in Mycobacterium tuberculosis. PLoS Pathog 21, e1013207. 10.1371/journal.ppat.1013207.

47. Bendre, A.D., Peters, P.J., and Kumar, J. (2021). Recent Insights into the Structure and Function of Mycobacterial Membrane Proteins Facilitated by Cryo-EM. J Membr Biol 254, 321–341. 10.1007/s00232-021-00179-w.

48. Kurz, S.G., and Rivas-Santiago, B. (2020). Time to Expand the Picture of Mycobacterial Lipids: Spotlight on Nontuberculous Mycobacteria. Am J Respir Cell Mol Biol 62, 275–276. 10.1165/rcmb.2019-0324ED.

49. Tran, T., Bonham, A.J., Chan, E.D., and Honda, J.R. (2019). A paucity of knowledge regarding nontuberculous mycobacterial lipids compared to the tubercle bacillus. Tuberculosis 115, 96– 107. 10.1016/j.tube.2019.02.008.

50. Poulton, N.C., Azadian, Z.A., DeJesus, M.A., and Rock, J.M. (2022). Mutations in rv0678 Confer Low-Level Resistance to Benzothiazinone DprE1 Inhibitors in Mycobacterium tuberculosis. Antimicrob Agents Chemother 66. 10.1128/AAC.00904-22.

51. Snobre, J., Villellas, M.C., Coeck, N., Mulders, W., Tzfadia, O., de Jong, B.C., Andries, K., and Rigouts, L. (2023). Bedaquiline- and clofazimine-selected Mycobacterium tuberculosis mutants: further insights on resistance driven largely by Rv0678. Sci Rep 13. 10.1038/S41598-023-36955-Y.

52. Huang, L., Matsuo, M., Calderón, C., Fan, S.H., Ammanath, A.V., Fu, X., Li, N., Luqman, A., Ullrich, M., Herrmann, F., et al. (2022). Molecular Basis of Rhodomyrtone Resistance in Staphylococcus aureus. mBio 13, e03833–21. 10.1128/MBIO.03833-21.

53. Horan, K.L., Freeman, R., Weigel, K., Semret, M., Pfaller, S., Covert, T.C., van Soolingen, D., Leão, S.C., Behr, M.A., and Cangelosi, G.A. (2006). Isolation of the genome sequence strain Mycobacterium avium 104 from multiple patients over a 17-year period. J Clin Microbiol 44, 783–789. 10.1128/JCM.44.3.783-789.2006.

54. Quach, D., Sharp, M., Ahmed, S., Ames, L., Bhagwat, A., Deshpande, A., Parish, T., Pogliano, J., and Sugie, J. (2025). Deep learning–driven bacterial cytological profiling to determine antimicrobial mechanisms in Mycobacterium tuberculosis. Proceedings of the National Academy of Sciences 122. 10.1073/pnas.2419813122.

55. Schindelin, J., Arganda-Carreras, I., Frise, E., Kaynig, V., Longair, M., Pietzsch, T., Preibisch, S., Rueden, C., Saalfeld, S., Schmid, B., et al. (2012). Fiji: an open-source platform for biological-image analysis. Nat Methods 9, 676–682. 10.1038/nmeth.2019.

56. Ducret, A., Quardokus, E.M., and Brun, Y. V. (2016). MicrobeJ, a tool for high throughput bacterial cell detection and quantitative analysis. Nat Microbiol 1, 16077. 10.1038/nmicrobiol.2016.77.

57. Bosch, B., DeJesus, M.A., Poulton, N.C., Zhang, W., Engelhart, C.A., Zaveri, A., Lavalette, S., Ruecker, N., Trujillo, C., Wallach, J.B., et al. (2021). Genome-wide gene expression tuning reveals diverse vulnerabilities of M. tuberculosis. Cell 184, 4579-4592.e24. 10.1016/j.cell.2021.06.033.

58. Chengalroyen, M.D., Mehaffy, C., Lucas, M., Bauer, N., Raphela, M.L., Oketade, N., Warner, D.F., Lewinsohn, D.A., Lewinsohn, D.M., Dobos, K.M., et al. (2024). Modulation of riboflavin biosynthesis and utilization in mycobacteria. Microbiol Spectr 12. 10.1128/spectrum.03207-23.

59. Rock, J.M., Hopkins, F.F., Chavez, A., Diallo, M., Chase, M.R., Gerrick, E.R., Pritchard, J.R., Church, G.M., Rubin, E.J., Sassetti, C.M., et al. (2017). Programmable transcriptional repression in mycobacteria using an orthogonal CRISPR interference platform. Nature Microbiology 2017 2:4 2, 1–9. 10.1038/nmicrobiol.2016.274.

