## Supplementary Figures and Tables for "Chlorotonils exhibit potent activity against *Mycobacterium tuberculosis*, while resistance is mediated by MmpR5-MmpL5"

***Table SI 1: Tabulated summary of all reference compounds used in BCP with corresponding similarity score to DHG.***

| Similarity score | Comparator | MoA |
|---|---|---|
| 87 | Bedaquiline | ATP Synthase |
| 86 | CCCP | Proton Ionophore |
| 71 | Q203 | Cytochrome bcc complex |
| 23 | Cycloserine | Cell Wall (peptidoglycan) |
| 25 | Meropenem | Cell Wall (peptidoglycan) |
| 22 | AU1235 | Mmpl3 inhibitor |
| 11 | Ethambutol | Cell Wall (arabinogalactan) |
| 19 | TBA-7371 | Cell Wall (arabinans synthesis) |
| 33 | SQ109 | MmpL3 Inhibitor |
| 5 | Gatifloxacin | DNA Gyrase |
| 22 | Mitomycin_C | DNA replication |
| 14 | Novobiocin | DNA replication |
| 35 | Cerulenin | Fatty acid synthesis |
| 27 | Ethionamide | Mycolic acid synthesis |
| 36 | Isoniazid | Mycolic acid synthesis |
| 47 | Pretomanid | Mycolic acid synthesis |
| 61 | Nisin | Membrane active |
| 37 | Thioridazine | Membrane active (undefined) |
| 24 | Actinomycin_D | Transcription |
| 22 | Rifabutin | Transcription |
| 37 | Clarithromycin | Translation (macrolide) |
| 23 | Gentamicin | Translation (aminoglycoside) |
| 10 | Sutezolid | Translation (oxazolidinone) |
| 61 | DMSO | DMSO |

***Table SI 2: All sensitizers (left) and resistors (right) identified by the CRISPRi screen using DHG at 0.5xMIC against M. tuberculosis.***

| gene | Short Description | change | gene | Short Description | change |
|---|---|---|---|---|---|
| **Rv1109c** | Cell division regulation | -3.39 | **Rv0678** | Efflux pump regulator | 2.38 |
| **mmpS5** | Efflux pump partner | -3.12 | **kasB** | Mycolic acid synthesis | 1.68 |
| **mmpL5** | Efflux pump transporter | -2.86 | **fabD** | Fatty acid synthesis | 1.61 |
| **Rv0479c** | Oxidoreductase stress response | -2.63 | **guaB3** | GMP synthesis enzyme | 1.44 |
| **Rv1830** | Transcriptional regulator | -2.17 | **Rv2970A** | Polyketide synthesis enzyme | 1.41 |
| **Rv3802c** | Lipid metabolism enzyme | -2.14 | **kasA** | Mycolic acid synthesis | 1.38 |
| **dnaA** | Initiates DNA replication | -1.79 | **secA1** | Protein translocation ATPase | 1.32 |
| **ppk** | Polyphosphate synthesis enzyme | -1.66 | **acpP** | Acyl carrier protein | 1.30 |
| **Rv1222** | Transcription termination factor | -1.59 | **sigE** | Stress response sigma factor | 1.27 |
| **Rv1218c** | Multidrug efflux pump | -1.53 | **accD6** | Fatty acid carboxylase | 1.21 |
| **arsA** | Arsenic resistance ATPase | -1.53 | **Rv1960c** | ABC transporter component | 1.15 |
| **Rv1219c** | Transcriptional repressor | -1.42 | **phoP** | Two-component regulator | 1.13 |
| **Rv1217c** | ABC transporter component | -1.40 | **Rv3764c** | ABC transporter component | 1.12 |
| **dnaB** | DNA helicase replication | -1.33 | **pcaA** | Mycolic acid cyclopropane | 1.12 |
| **Rv0204c** | PE-PGRS protein | -1.31 | **fadE5** | Fatty acid degradation | 1.08 |
| **Rv1707** | Transcriptional regulator | -1.30 | **accA3** | Acetyl-CoA carboxylase | 1.05 |
| **ideR** | Iron-dependent regulator | -1.19 | **Rv3651** | Oxidoreductase activity | 1.04 |
| **rne** | Ribonuclease E | -1.16 | **fadD32** | Mycolic acid activation | 1.04 |
| **galU** | UDP-glucose synthesis | -1.11 | **Rv2971** | Polyketide synthesis | 1.02 |
| **rpmG** | Ribosomal protein L33 | -1.09 | | | |
| **whiA** | Cell division regulator | -1.04 | | | |
| **Rv1906c** | Stress response protein | -1.02 | | | |

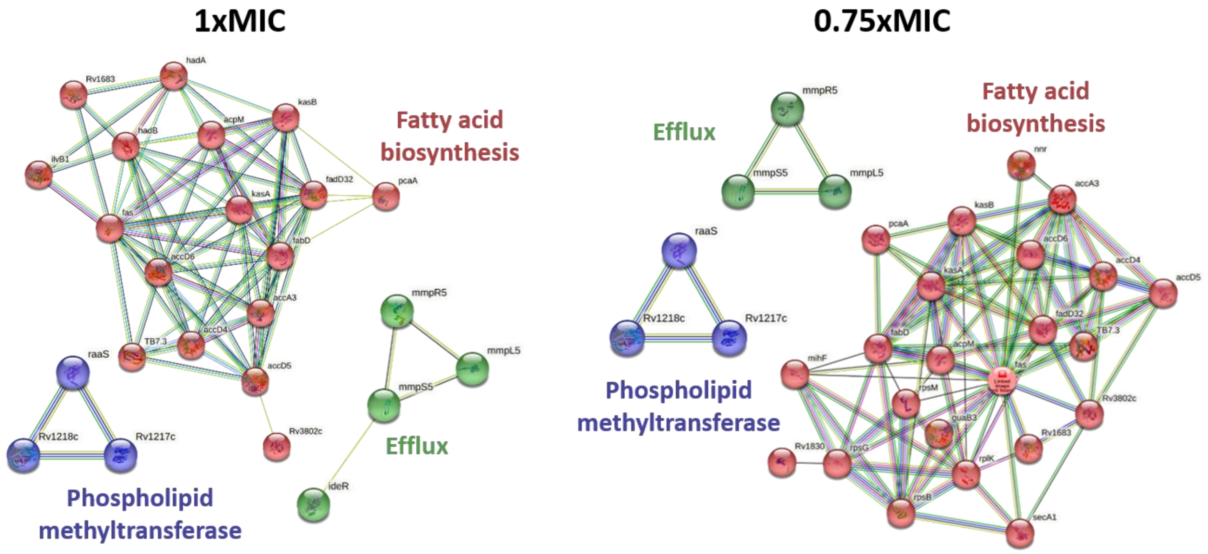


***Figure SI 1: STRING network analyses of significant CRISPRi hits reveal efflux, fatty acid biosynthesis and phospholipid methyl transferase clusters functionally enriched.***

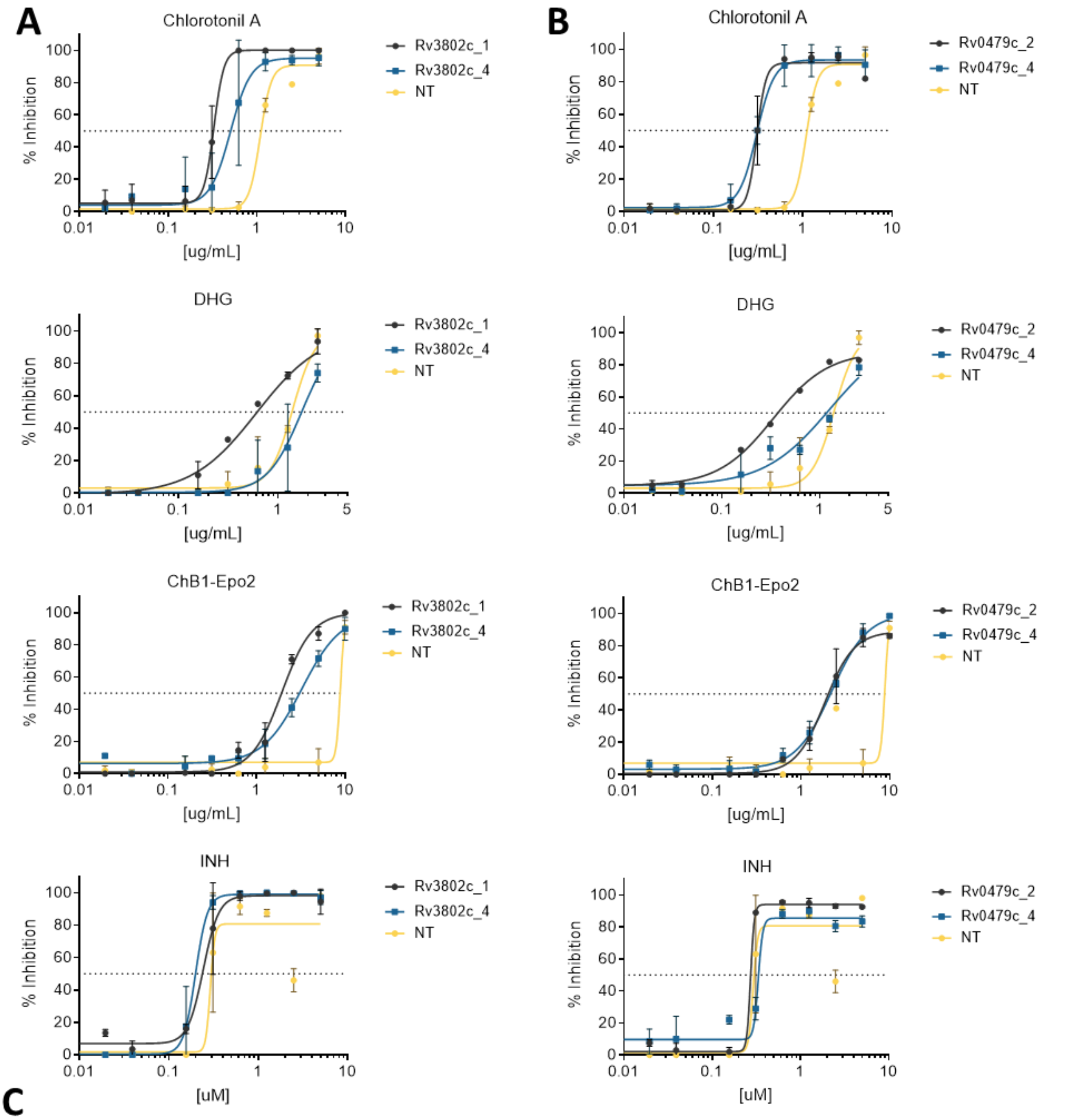


**C**

| Drug | non-induced $IC_{50}$ [µM] | Rv3802c | | | | Rv0479c | | | |
|---|---|---|---|---|---|---|---|---|---|
| | | $IC_{50}$ [µM] | | fold-change | | $IC_{50}$ [µM] | | fold-change | |
| | | strong sgRNA | weak sgRNA | strong sgRNA | weak sgRNA | strong sgRNA | weak sgRNA | strong sgRNA | weak sgRNA |
| ChA | 1.092 | **0.329** | **0.505** | **3.32** | **2.16** | **0.306** | **0.306** | **3.56** | **3.57** |
| DHG | 1.376 | **0.616** | 1.724 | **2.23** | 0.80 | **0.311** | 1.167 | **4.43** | 1.18 |
| BE | 8.746 | **1.916** | **3.030** | **4.56** | **2.89** | **1.849** | **2.146** | **4.73** | **4.08** |
| INH | 0.291 | 0.243 | 0.199 | 1.20 | 1.46 | 0.273 | 0.333 | 1.07 | 0.87 |

***Figure SI 2: Moderate hypersensitization of both Rv3802c and Rv0479c towards chlorotonils upon depletion.*** *Rv3802c (A) or Rv0479c (B) was gradually depleted (3-day ATc predepletion) using either a potent sgRNA (Rv3802-1 or Rv0479_2), or a weak sgRNA (Rv3802-4 or Rv0479-4). Treatment was performed using natural product Chlorotonil A, DHG, ChB1-Epo2 (BE) and isoniazid (INH), the off-target drug control at different concentrations (n=2, mean and SD is shown). Data were fitted using non-linear regression in GraphPad Prism 10.5. C) Quantitative evaluation of A and B. Data is expressed as $IC_{50}$ or as fold-change in relation to the non-target (NT) sgRNA strain.*

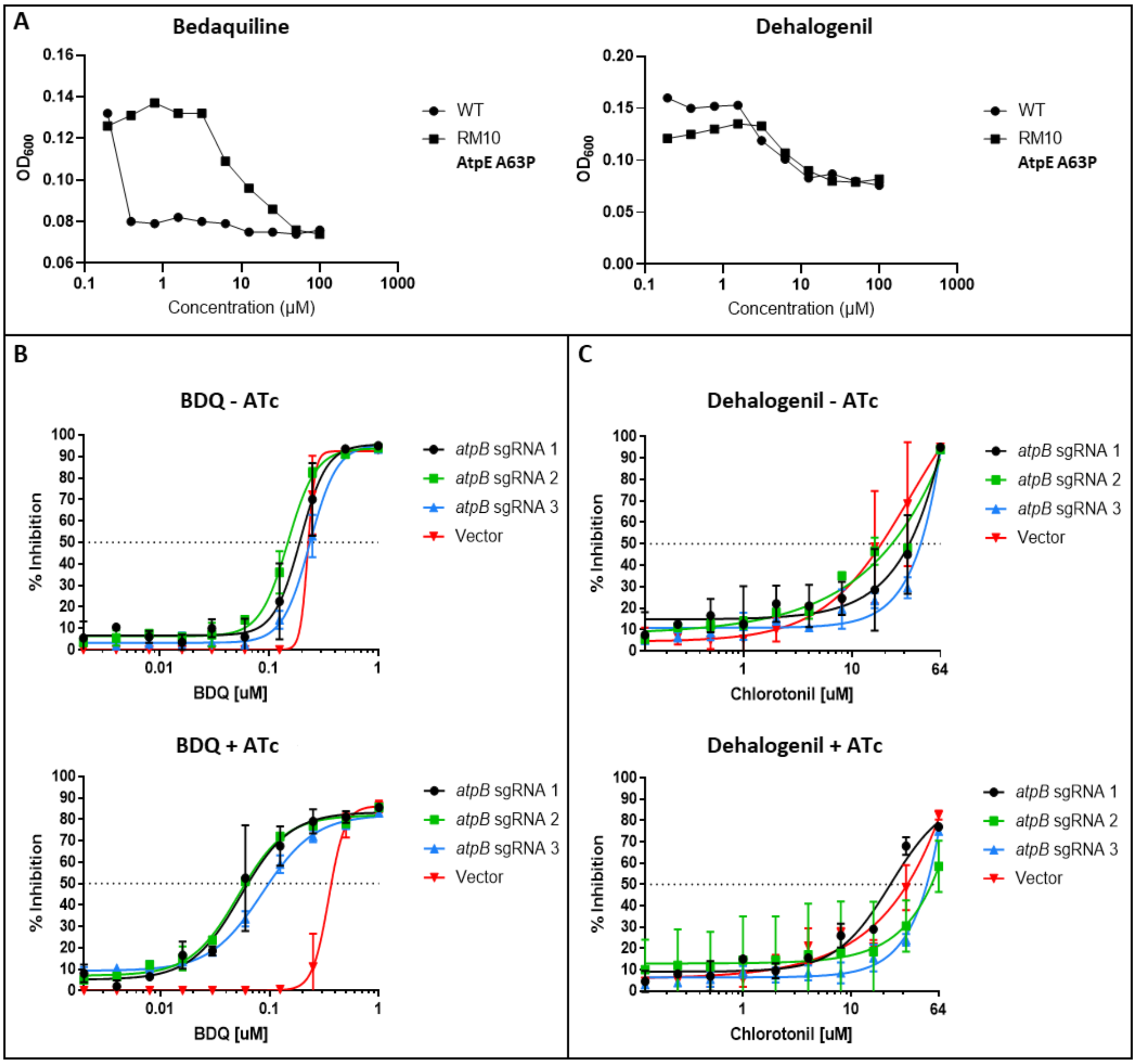


***Figure SI 3: Dehalogenil does not display on-target activity against AtpE or AtpB.*** *A) Mtb RM10 carrying mutations in atpE is less susceptible to BDQ but not DHG. B/C) AtpB hypomorphs are more susceptible to BDQ but not to DHG. A panel of atpB hypomorphs was tested, namely a potent (1), medium (3), or weak sgRNA (2) targeting strain. Treatment was performed using DHG or BDQ at different concentrations (n=2, mean and SD is shown).*
